# DynMoCo: a Novel AI Framework to Reveal Modular Substructures of Protein From Molecular Dynamics

**DOI:** 10.64898/2026.02.08.704355

**Authors:** Lingchao Mao, Mingu Kwak, Amir Hossein Kazemipour Ashkezari, Zhenhai Li, Yunfeng Chen, Peiwen Cong, Jung Hun Phee, Sooyeon Kang, Jing Li, Cheng Zhu

## Abstract

Proteins are dynamic molecular machines whose functions are determined by their structures. While static structures can offer initial insights or hypotheses about protein function, they are often insufficient for a detailed mechanistic understanding. Molecular dynamics (MD) simulations provide atomistic view of protein’s dynamic motion and conformational change, but the resulting high-dimensional data are challenging to interpret. Traditional summary statistics and dimensionality-reduction methods often focus on global motions and can overlook regional, yet functionally critical motions. Inspired by approaches from social network science, we introduce a novel perspective for analyzing MD simulations through dynamic community detection, where molecules are modeled as time-evolving graphs, and communities of residues or atoms that move coherently or exhibit functional coupling are identified. We present DynMoCo, a novel deep learning framework that integrates graph convolutional networks with recurrent models for end-to-end dynamic community detection on molecular graphs. Given a MD trajectory, DynMoCo identifies spatially grounded substructures, tracks their evolution over time, and can incorporate structural knowledge to ensure physically meaningful communities. We provide a library of custom-written scripts to allow users to extract and visualize these communites on the MD simulated molecules in motion. We demonstrate the method on force-ramp and force-clamp steered MD simulations of three integrin systems, revealing modular substructures within known domains and characterizing their conformational rearrangements during force-induced unbending. By reducing high-dimensional MD data into interpretable communities, this approach offers new insights into the intrinsic organization and dynamic function of complex biomolecular systems.

**SIGNIFICANCE:** Proteins often perform their functions through dynamic, locally coordinated motions. Molecular dynamics simulations provide detailed views of these motions but produce high-dimensional data that are challenging to analyze and interpret. We present a novel deep learning model that analyzes molecular dynamics simulations data and identifies structurally coherent and potentially functionally related communities, while tracking their temporal evolution. This analysis tool provides a novel way to analyze MD data transforming it into interpretable representations of modular dynamic, enabling discovery of new mechanistic insights and advancing our understanding of how molecular motions drive biological function.

## INTRODUCTION

The principle that function follows structure (structure–function relationship) is foundational in biology, spanning across length scales from whole organisms down to molecular machines such as proteins. This relationship underpins the field of structural biology, where solving the three-dimensional (3D) structures of biological macromolecules is central to understanding their function. Recent advances in cryogenic electron microscopy (cryo-EM) and artificial intelligence (AI)-based structure prediction have dramatically expanded our ability to obtain high-quality protein structures, both experimentally and computationally.

Although static structures provide some working hypotheses for protein functions, such structural information is usually insufficient for a full understanding of function. The reason is multi-fold. The protein structures solved, regardless of whether they were done experimentally or computationally, are static, reside at the lowest energy state at zero degree absolute temperature, and are free of applied force. Under physiological conditions, by comparison, proteins are dynamic as they function at body temperature, may assume multiple conformational states and transition among them, and may be subjected to mechanical forces that modulate their conformations and induce conformational changes. Also, the protein structures are information-rich, as they provide 3D coordinates of all atoms of the protein in space, yet it is often not easy or even impossible to translate such information into understanding and knowledge.

Molecular dynamics (MD) simulations have the potential to address this gap by modeling proteins in solvated, thermally equilibrated environments and generating time-resolved trajectories at atomic resolution. MD has become a powerful tool for studying protein folding, allostery, ligand binding, and mechanically induced conformational changes (1). By simulating the thousands to millions of atoms in motion at femtosecond resolution, MD provides a virtual nanoscope into biomolecular dynamics and a physics-based framework for generating mechanistic hypotheses about biological function. Yet despite our ability to generate massive amounts of MD data, our ability to comprehend them and utilize them to understand protein function is disappointingly limited. MD trajectories are high-dimensional, continuous, unlabeled, and noisy, and although atomic motions are governed by well-defined physical interactions, their collective behavior is often difficult to interpret directly.

Traditional MD analysis relies heavily on hand-crafted features such as Cartesian coordinates, interatomic distances, contact maps, or dihedral angles. While effective for specific systems or hypotheses, these approaches are often system-dependent and may miss unexpected or emergent mechanisms. This challenge has led to a shift toward advanced analytical methods, particularly graph-based network analysis and machine learning (2). Graph representation where residues or atoms are nodes and their interactions define edges (3) naturally encode molecular topology and spatial organization. This formulation shifts the focus from individual motions to cooperative patterns of interaction, offering a principled way to study collective behavior in MD trajectories. To date, most successful applications of machine learning and deep learning in molecular science have focused on static structures, including protein function prediction, drug discovery, and protein design (4). In contrast, learning directly from dynamic molecular data remains comparatively underexplored.

A growing body of work has emphasized that proteins operate as dynamic conformational ensembles, and that allosteric regulation arises from network-based propagation of structural perturbations rather than from static structures alone. Prior approaches—including Dynamic Cross Correlation Matrices (DCCM) (5), Linear Mutual Information (LMI) (6), perturbation response scanning, evolutionary trace analysis, and machine learning–based residue response maps—have provided powerful means of identifying correlated motions or allosteric communication pathways, but these methods operate on static or time averaged graphs and therefore cannot capture the temporal evolution of mesoscale organizational units during conformational transitions. As highlighted in the comprehensive review by Nussinov and colleagues (7), allosteric networks involve time dependent changes in residue–residue connectivity, reflecting shifts in ensemble populations, pathway usage, and signaling bias. However, current network based approaches lack mechanisms to maintain community identity across frames, making them unable to follow key events such as community emergence, disappearance, splitting, or merging during structural rearrangements.

While community detection is well established in social and transportation analysis (8, 9), its application to dynamic molecular graphs still face unresolved challenges. Most dynamic community detection methods apply static algorithms independently to each snapshot and then attempt to align communities across time (10), or impose temporal smoothness constraints based on previous assignments (11, 12). These strategies may fail to capture complex, non-smooth conformational transitions characteristic of some biomolecular systems like the one studied in this work. Second, most existing network analysis methods rely primarily on network topology and ignore rich node- and edge-level attributes that are critical for molecular interpretation (8, 13). Third, matrix factorization and evolutionary clustering approaches are often limited by linear assumptions or predefined temporal regularization, which restricts their ability to model nonlinear and heterogeneous dynamics (14, 15). Recent deep learning methods have more flexible learning-based frameworks with differentiable clustering objectives (16, 17). However, there remain very few unsupervised models that can jointly learn dynamic representations and community structure in an end-to-end manner (18), and none are designed specifically for molecular graphs.

In this work, we introduce **DynMoCo**, the first end to end deep learning framework for dynamic community detection on molecular graphs. By integrating graph convolutional networks (GCN) with recurrent temporal modeling (RNN) from deep learning, DynMoCo identifies spatially grounded substructures and robustly tracks their evolution throughout an MD trajectory, yielding communities that remain coherent across time and align closely with known structural and functional domains. Unlike static community detection methods, which must be applied independently to each snapshot and suffer from label switching and loss of temporal interpretability, DynMoCo learns a temporally consistent representation of modular organization, enabling quantitative and visual analyses of how substructures reorganize during force induced or thermally driven conformational change. In addition, DynMoCo incorporates topological priors through a knowledge-informed modularity loss to identify structurally and functionally meaningful communities. This positions DynMoCo not only as a methodological advance in MD analysis but also as a tool that provides a new perspective to analyze protein dynamics and reorganization.

## MATERIALS AND METHODS

### Molecular graph construction and sampling

MD simulations track the physical movements of a molecular system comparising of *N* atoms over time and generate a time-resolved trajectory of three-dimensional atomic positions, which can be transformed into a sequence of molecular interaction graphs. Each MD simulation is represented as a sequence of graphs *G*_*t*_ = (*V, E*_*t*_), where the node set *V* = {*v*_1_, *v*_2_, …, *v* _*N*_}, |*V*| = *N* corresponds to residues of the molecular system, and the edge set *E*_*t*_ ⊆ *V* × *V*, |*E*_*t*_| = *M*_*t*_ represent meaningful pairwise interactions or contacts between residues at time *t*. We assume a fixed node set across time, while edges may appear or disappear. Each node is associated with a feature vector encoding geometric (e.g. coordinates) or physicochemical properties (e.g. mass, electronegativity), denoted as *X*_*t*_ = ℝ ^*N* ×*D*^. An edge is defined based on existence of covalent bonds, which remain largely stable throughout the conformational change, and non-covalent interactions. There are four categories of non-covalent interactions (hydrogen bonds, ionic interactions, van der Waals forces, and hydrophobic interactions). For simplicity, we determine non-covalent interactions based on a predefined distance cutoff and treat the four categories of non-covalent interactions the same, although more sophisticated algorithms or software packages can be used to determine each type of non-covalent interactions. In cases one residue concurrently interacts with more than one other residue with cross-talk, the residue will have multiple edges connecting it with other residues.

In an MD simulation with high spatiotemporal resolution, successive frames may differ only by high frequency, stationary fluctuations. Modeling every frame is computationally unnecessary and may introduce noise. Instead, we employ a principled sampling strategy to construct graphs based on a subset of sampled frames. MD simulations can be broadly categorized into two types: (i) free MD, in which the molecule evolves in the solvent under equilibrium without external perturbations, and steered MD (SMD), where external forces are applied to simulate biologically relevant processes such as ligand binding, protein unfolding, or mechanical stress. SMD can be further divided into force-clamp SMD, where the molecule is subjected to a certain extension under constant force, and force-ramp SMD, where the molecule is pulled at constant velocity through a linear spring to generate a ramp force.

We proposed the following frame sampling strategy to construct the molecular graph for different types of MD simulations:

#### Free or force-clamp SMD

the molecular system undergoes dynamic fluctuations around a fixed (intermediate) state and hence, we assume the frames, sampled at sufficiently low frequency across the trajectory, are independent and identically distributed observations of the state. To identify statistically meaningful contacts within the state, we compute pairwise distances between the C*α* atoms of residue pairs in each frame, followed by a one-sided one-sample t-test to assess whether the average distance is significantly less than 5Å. Bonferroni correction is applied to account for multiple comparisons, yielding a robust contact graph that reflects persistent interactions. Covalent bonds are deterministically included as edges based on the molecular topology.

#### Force-ramp SMD

the molecule is pulled by an increasing force leading to force-induced conformational changes such as unbending. Clearly, the subsequent frames are no longer independent and identically distributed. Instead, we extract non-overlapping segments from the trajectory, each time interval representing an intermediate state of interest. Within each segment, we obtain bootstrapped samples of frames and apply the same statistical test described above to construct a molecular graph. The number and location of the segments, the number of frames and bootstrap samples can be tailored to the specific dynamics of the system under study, balancing temporal resolution with computational efficiency.

This preprocessing yields a sequence of graphs with statistically significant edges to serve as robust inputs for dynamic community detection.

### Dynamic Molecular Community Detection: DynMoCo

Given a temporal graph sequence 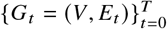, our objective is to detect dynamic communities 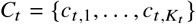, for each time point *t* ∈ { 0,…, *T*} . Each community is a group of nodes that (i) are densely interconnected in the molecular graph and (ii) exhibit similar node attributes, reflecting cohesive structural or functional modules. We aim to track the formation, evolution, and dissolution of these communities over time, enabling interpretable insights into the modular reorganization of biomolecules during dynamic processes such as conformational transitions or mechanical unfolding. After generating the algorithm outputs, residues assigned to the same community can be color-coded and visualized on the protein structure during conformational dynamics using the user-controlled molecular visualization software VMD (19).

In this work, we develop DynMoCo, a novel deep learning framework designed to model time-evolving communities from a sequence of molecular graphs. DynMoCo combines the flexibility and representation power of graph neural networks with the temporal modelling capabilities of recurrent neural networks to perform end-to-end dynamic community detection. This model outputs soft node-level community assignments 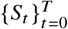, which can be converted into hard assignment labels by taking the one with maximum probability. DynMoCo’s architecture comprises two main components: a graph convolutional encoder for node embedding and a recurrent neural network for community assignment evolution.

### Node embedding via graph convolution

As molecules can be naturally represented as graphs, we apply a Graph Convolutional Network (GCN) as the encoder to extract expressive representations of nodes at each time point. GCNs can advantageously integrate both structural information (i.e., graph topology) and node-level attributes (i.e., residue properties) to generate node embeddings.

At time *t*, a single-layer GCN maps the node features into higher-dimensional embeddings as follows:

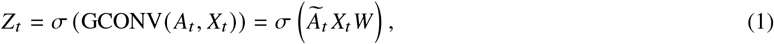

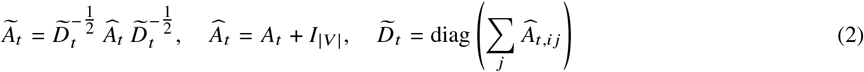

where *Z*_*t*_ ∈ ℝ^*N* × *H*^ is the node embedding matrix, *W* ∈ ℝ^*D* × *H*^ is the learnable weight matrix, *A*_*t*_ ∈ℝ^*N* × *N*^ is the adjacency matrix where 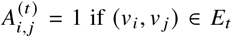 and 0 otherwise, Ã_*t*_ is the renormalized adjacency matrix, and σ is the activation function. We used SeLU activation function (20) instead of ReLU for better convergence.

The neighborhood aggregation mechanism of GCNs allows each node’s embedding to be computed by its local graph structure (i.e., its neighbors), which serves as a denoising effect of the high frequency fluctuations across frames and allows capturing potential functional relationships among neighboring residues. Stacking multiple GCN layers expands the receptive field, enabling the model to integrate information across larger molecular neighborhoods.

### Temporal modeling via recurrent neural networks

When a molecule undergoes force-induced conformational change, its underlying substructures are expected to reorganize based on local interactions. These force-driven dynamics and rearrangements can be central to the molecule’s function. Hence, we design the deep learning model to detect structural dynamic evolution of the communities.

To capture temporal evolution of communities, we use a Gated Recurrent Unit (GRU) to learn the community assignment matrix *S*_*t*_ ∈ ℝ^*N* ×*C*^. GRU is a type of recurrent neural network (RNN) designed to model sequential data by using “gates” to control the flow of information and mitigate the problems of vanishing and exploding gradients of standard RNNs. We use GRU to learn the evolution of node-community affiliations over time, taking into account both newly observed node embeddings and prior community memberships:

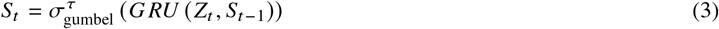

where 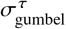 is the Gumbel-Softmax activation function (21). The temperature parameter τ > 0 controls the sharpness of the distribution with lower values of τ producing outputs closer to one-hot vectors (i.e. hard assignments), while higher values of τ yields softer, more distributed community assignments. We adopt Gumbel-Softmax to generate differentiable yet approximately discrete community assignments that are more suitable for modularity computation, which will be discussed in detail in the next section.

A standard GRU layer is defined as follows:

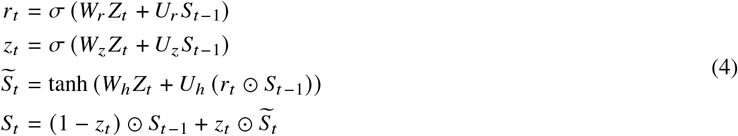

where σ is sigmoid, *r*_*t*_, *z*_*t*_ are the reset and update gates, *S*_*t*_ is a hidden state candidate, *W*_*r*_, *W*_*z*_, *W*_*h*_, *U*_*r*_, *U*_*z*_, *U*_*h*_ are learnable weights, and ⊙ denote element-wise multiplication. These gates regulate the flow of temporal information, allowing the model to incorporate new information while selectively retaining meaningful past information. The hidden state, i.e. the community assignments, evolves gradually unless there is consistent evidence of a shift. This makes GRUs robust to noisy perturbations in MD data while capturing long-term dynamics.

Proper initialization of the hidden state, *S*_0_, is critical for stable training and convergence. To this end, we employ the Louvain algorithm (8), an edge-based heuristic for community detection on static graphs, to compute the initial community assignments on *G*_0_ as the initialization of GRU’s hidden state.

### Community pooling and embedding

Once the community assignment matrix *S*_*t*_ is learnt, we perform graph pooling to obtain higher-level community embeddings, *H*_*t*_ and coarsened adjacency matrix,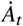:

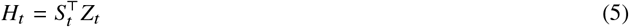

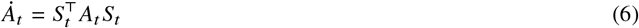

### Unsupervised Learning with Knowledge-Informed Modularity Loss

#### Modularity optimization

To identify structurally consistent communities in molecular systems, we adopt an unsupervised objective based on modularity maximization. Modularity is a commonly used metric that quantifies the quality of division of a graph into communities, comparing the actual edge density within clusters to that expected in a random graph with the same node degree distribution. Formally, the modularity (*Q*) is defined as:

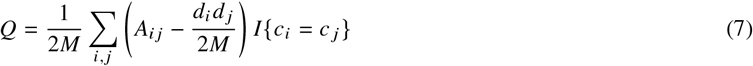

where *d*_*i*_ = Σ_*j*_ *A*_*i j*_ is the degree of node *v*_*i*_, *c*_*i*_ is the community to which node *v*_*i*_ is assigned, *M* is the total number of edges in the graph, *I* {*c*_*i*_ = *c* _*j*_{ is an indicator function of whether *v*_*i*_ and *v* _*j*_ belong to the same community. Briefly, 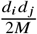 is the expected number of edges between nodes and *A*_*i j*_ is the observed number of edges. *Q* ranges from − 1 to 1, and a larger *Q* leads to a better community partition.

Modularity maximization can be written as a constrained optimization of the following form:

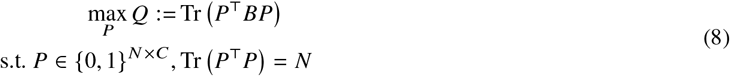

where 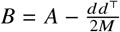 is the modularity matrix, and *P* = { *p*_1_, …, *p*_*N*_} ∈ {0, 1}^*N* ×*C*^ is a one-hot matrix indicating community assignment of each node in the graph.

Modularity maximization encourages densely connected groups of nodes with sparse connections between groups and is a popular approach in community detection. However, maximizing modularity is proven to be NP-hard (22). The optimization constraints can be relaxed to define a differentiable modularity loss as:

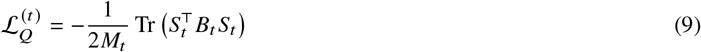

Directly optimizing this objective may lead to modularity collapse, where all nodes are assigned to the same cluster yielding a trivial solution that would trap gradient-based optimizers. To avoid collapse, we adopt the strategy used in Deep Modularity Networks (DMoN) (23), which introduces a regularizer based on the Frobenius norm of soft cluster membership counts:

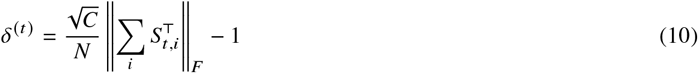

The regularized relaxed modularity loss becomes:

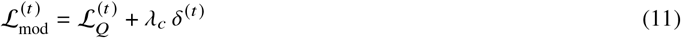

where *λ*_*c*_ is a hyper-parameter that controls the strength of the regularizer.

#### Knowledge consistency

For some commonly-studied molecular systems, prior structural knowledge such as domains is often available and can provide valuable information to weakly supervise model learning. Domains are groups of spatially or functionally related residues that form a compact region within a protein’s polypeptide chain and possesses a stable three-dimensional structure, and often performs a specific function, such as binding to another molecule or catalyzing a chemical reaction. Hence, domains form a higher-level modular structure that communities should respect. While a domain usually contain multiple communities, communities generally do not span across multiple domains.

To incorporate this structural knowledge, we first define a domain co-membership mask *D* ∈ {0, 1}^*N* ×*N*^ as:

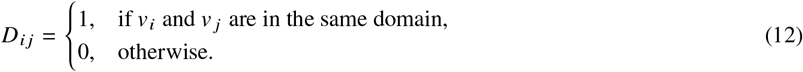

We then introduce a knowledge consistency loss that penalizes residues from different domains having similar soft community assignments:

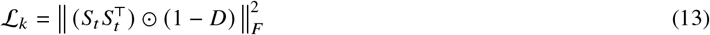

This loss is unidirectional as it penalizes cross-domain similarity of communities but allows for multiple communities to be formed within a single domain.

#### Sparse structural change regularization

Since communities represent substructures of a molecular system that are functionally and structurally related, we assume most communities exhibit temporal stability unless there is strong evidence of community structure change. In other words, structural changes such as splitting, merging, appearance, or disappearance of communities should be rare. To enforce this, we introduce a temporal smoothness regularization that penalizes large differences in community assignments across adjacent time steps:

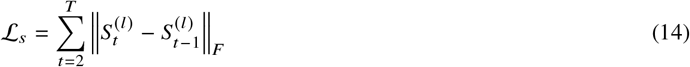

#### Final objective

The final unsupervised loss function combines the modularity loss, knowledge consistency loss, and temporal sparsity loss as:

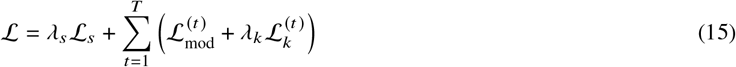

where *λ*_*s*_, *λ*_*k*_ are hyper-parameters that control the strength of sparsity regularizer and the influence of domain knowledge.

This framework enables end-to-end unsupervised dynamic community detection from any given graph sequence, combining the GCN’s strength at leveraging both node features and network topology for graph representation learning with the RNN’s capability at temporal modeling of complex long sequences. The integration of structural knowledge, if available, can further weakly supervise the learning process to find physically-meaningful communities on sparse, noisy molecular graphs.

### PageRank Analysis Identifies Mechanically Responsive Residues

The community detection algorithm provides a community-level characterization of molecular dynamics. To complement these results with finer-grained identification of residues critical for force propagation or structural rearrangements, we applied the PageRank algorithm (24) to the molecular graphs. Originally developed to rank webpages, PageRank (24) identifies nodes that are central or influential in a network by considering both the number of incoming edges and the importance of the linking nodes. Although PageRank is no longer the sole algorithm used by Google to rank search results, it remains one of the most widely recognized and influential methods for identifying central nodes in complex networks (25).

In molecular graphs, we adapt PageRank (25) to highlight residues that mediate interactions and transmit conformational changes. Nodes with many incoming edges, particularly from highly ranked nodes, achieve higher scores; thus, even a residue with few but strategically important contacts may rank higher than one with numerous weak connections. To reflect structural dynamics, we weight edges by residue–residue pairwise distance changes. Under this formulation, residues that are both well-connected and strategically positioned within structural pathways, while undergoing substantial positional shifts relative to their neighbors, achieve high PageRank scores. This network-based metric identifies residues that are topologically central and undergo large distance changes, serving as a proxy for residues that participate in mechanically responsive pathways during conformational change.

To construct the input graph for PageRank, each node represents a residue, and an edge is added between two residues if: (i) they are in statistically significant contact (mean distance < 5Å, one-sided t-test with Bonferroni correction) at the start or end of the conformational change, and (ii) their inter-residue distance changes significantly over time. Edges are weighted by the normalized magnitude of distance change. PageRank applied to this weighted graph yields residue-level scores that complement the community-level analysis by quantifying mechanical importance within conformational transitions. We explicitly note that, the PageRank score is not derived from a direct force measurement (e.g., from force distribution analysis), but instead reflects a residue’s role as a mechanically key hub of conformational change by integrating both local interactions and the global network context.

### Molecular Dynamics Simulations

To demonstrate the versatility of the model, we applied DynMoCo to analyze MD datasets of three structurally related yet functionally distinct molecules of the integrin family generated. The simulation setups are described in the original publications of these datasets (26, 27) but are also be summarized briefly below.

#### Force-clamp SMD of *α*_5_*β*_1_ **and** *α*_*V*_*β*_3_

The simulations were performed in a previous study of some of us and analyzed resuts were published without including the raw data in the publication (26), which were used for the present study. The MD simulations were conducted using the ectodomain crystal structures of *α*_*V*_*β*_3_ (PDB: 3IJE) and *α*_5_*β*_1_ (PDB: 7NXD). GROMACS was used as the simulation engine. The systems were solvated using the TIP3P water model and ionized with Na^+^ and Cl^−^ to a physiological concentration of 150 mM. The CHARMM36 all-atom force field was used for proteins, with the CHARMM additive force field for glycans. Following energy minimization, annealing, and NVT/NPT equilibration with heavy-atom restraints, free MD and SMD simulations were performed. In the SMD phase, pulling forces were applied to the integrin head at 0.5 nm/ns while the C*α* atoms of the C-termini of the *α* and *β* chains were anchored. Force-ramp SMD simulations (with increasing head-to-tail tension) were performed to generate a conformational trajectory with increasing head-to-tail extensions. When the head-to-tail extensions reached given values—12, 14, 16, and 18 nm for *α*_5_*β*_1_ and 3, 11, 16, and 18 nm for *α*_*V*_*β*_3_—the pulling was halted and the intermediate structures were subject to further force-clamped SMD simulations (with constant head-to-tail tension) to capture a conformation ensemble under a constant force. In total, the dataset includes 5-6 replications of free MD simulations of bent integrins and clamped MD simulations at defined extension levels.

#### Force-ramp SMD of *α*_*V*_*β*_3_ **and** *α*_*IIb*_*β*_3_

This dataset was previously analyzed in a recent study (27) and was obtained from the publicly available link provided by the authors. Simulations were carried out using GROMACS with the CHARMM36 all-atom force field and TIP3P water model. Each integrin system was solvated in a cubic water box and ionized with 150 mM NaCl. After energy minimization via steepest descent and multi-phase NVT/NPT equilibration with gradually released backbone restraints, production simulations were conducted. Each integrin underwent 100 ns SMD simulations, during which a constant pulling force of 66.4 pN was applied to the ligand-binding site (residues in MIDAS, LIMBS, and ADMIDAS) to induce molecular extension, while transmembrane helices were constrained in the vertical axis to mimic membrane anchoring. Temperature was maintained at 310 K using a V-rescale thermostat, and pressure coupling was turned off during the production phase. Each integrin simulation had three independent replicates.

Table 1 summarizes the dataset characteristics. Movies displaying the MD simulations are provided in Videos S1-4. As the two force-clamp SMD datasets (*α*_5_*β*_1_ and *α*_*V*_*β*_3_) are simulated at four different extension levels representing four intermediate stages of the conformational change, we extracted four uniformly sampled segments (0–10, 63–73, 126–136, and 189–199 frames) from the force-ramp SMD simulations (*α*_*V*_*β*_3_ and *α*_*IIb*_*β*_3_) for simplicity of the comparison.

**Table 1:**
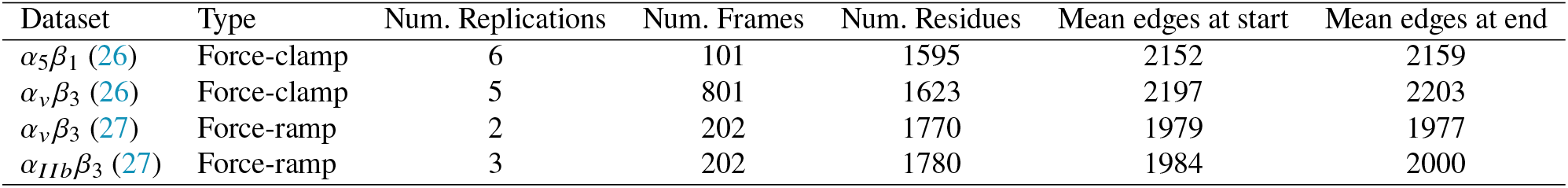
Summary characteristics of MD datasets analyzed in this study.

We use the proposed dynamic community detection model to analyze the four SMD datasets comparing with existing deep-learning based community detection methods. Model training details, complexity analysis, and competing methods are provided in Supplemental Materials.

#### Applying DynMoCo to other MD datasets

DynMoCo is designed as a general framework and is applicable to analysis of MD data for systems beyond integrins. No architectural modifications are required for different molecular systems. Adaptation primarily involves system-specific choices in graph construction and optional incorporation of prior knowledge.

Applying DynMoCo to a new system involves the following steps:

1. Dynamic graph construction: Given a SMD trajectory, define the intermediate states to be analyzed, including the temporal interval and sampling frequency, based on the dynamical characteristics of the biological system. Construct a sequence of graphs by defining nodes (e.g., residues or atoms) and edges (e.g. covalent and non-covalent interactions, pairwise distances).
2. Encoding structural priors: If available, incorporate prior knowledge such as known domains or functional regions by defining a comembership matrix to guide community detection. This step is optional and can be omitted for systems without well-characterized domain structures.
3. Running DynMoCo: Apply DynMoCo to the constructed temporal graph. The hyper-parameters *λ*_*s*_ and *λ*_*k*_ should be tuned for best performance. Users can refer to the provided Python implementation described in the Software section.
4. Community visualization and interpretation: Visualize the inferred dynamic communities using VMD and interpret their temporal evolution.

Note that DynMoCo is designed for analyzing SMD trajectories assuming the existence of temporally evolving community structures. For equilibrium simulations that fluctuate around a stable state without structural reconfiguration, static community detection methods such as Louvain or Girvan–Newman are more appropriate.

### Evaluation Metrics

We evaluated the performance of the models using widely used metrics for assessing the quality of graph partitions:

- **Modularity** measures the strength of division of a graph into communities by comparing the density of edges within communities to what would be expected in a random graph with the same degree distribution. Formally, modularity is defined in equation (7). It ranges from -0.5 to 1.0, where higher values (typically >0.3) indicate stronger community structure, and a value near 0 suggests no significant modular structure.
- **Conductance** quantifies the fraction of total edge volume that crosses the boundary of a community relative to the total number of edges incident to the community. For a community *S*, conductance is defined as

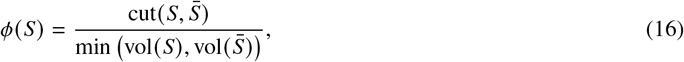

where cut 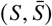 is the number of edges between *S* and its complement 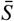, and vol (*S*) and vol 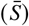 are the sum of degrees of nodes in *S* and 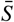, respectively. *ϕ* (*S*) ranges from 0 to 1, where lower values (closer to 0) reflect better-separated, more cohesive communities and a value near 1 indicates high inter-community connectivity.
- **Normalized Mutual Information (NMI)** evaluates the similarity between two community assignments by measuring the amount of shared information, normalized by their entropy. Given two partitions *X* and *Y*, NMI is defined as, where *I* (*X*; *Y*) is the mutual information between the partitions and H denotes entropy. NMI ranges from 0 (no mutual information) to 1 (perfect match).

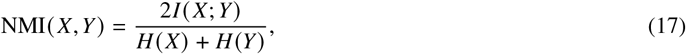
- **Adjusted Rand Index (ARI)** compares the similarity of two community partitions. Rand Index measures the agreement between two partitions by considering all pairs of samples and computing the proportion of pairs that are consistently assigned out of all possible pairs. ARI is rand index adjusted for chance, defined as:

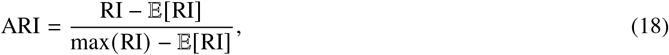

where 𝔼 [RI] is the expected value of rand index under random labelling. An ARI of 1 indicates perfect agreement, while 0 indicates random labeling.

Modularity and conductance were used to assess structural coherence and separability of the detected communities, while NMI and ARI were used to quantify community structure change over time and across molecules. Communities were also qualitatively assessment based on existing structural knowledge about the molecular systems.

### Software and Community Visualization

The code for running DynMoCo is publicly available in GitHub repository: https://github.com/lingchm/DynMoCo. For users interested in exploring community composition in detail, we provided custom scripts that load the community results into VMD software (19) and allow users to interactively visualize the molecular system overlaid with the communities with custom color-coding, representation format, community/residue/atom-level selections, and visualize community structure change.

## RESULTS AND DISCUSSION

### DynMoCo identifies structurally coherent and biologically relevant communities

#### DynMoCo outperforms existing benchmarks in community detection on dynamic graphs

The community detection performance for the force-clamp SMD datasets is summarized in Table 2 and for the force-ramp SMD datasets are shown in Table 3. Among competing methods, DMoN outperforms DEC due to its collapse regularization mechanism, though both struggle to generalize to dynamic settings. While DynAERNN is effective for temporal representation learning, its autoencoder-based, non-graph-based architecture, and two-stage pipeline limit the joint optimization of embeddings and community assignments. While MFC-TopoReg yielded more coherent communities than DynAERNN by its topological loss mechanism, the method is not designed for molecular graphs and yielded suboptimal results on MD data. In contrast, our proposed model DynMoCo consistently achieved better performance in both modularity and conductance metrics, demonstrating its ability to identify communities that are densely intra-connected and well-separated. While traditional community detection methods achieved high modularity scores at individual time points, they are designed for static community detection and exhibited poor alignment of community assignment over time, making community change events not interpretable (Table S11 and Figure S12). Qualitatively, DynMoCo recovers communities that are more aligned with known domains than the benchmark methods, showing its ability to discover biologically meaningful organization (Figure S1 and S13). Most communities identified by DynMoCo align closely with known protein domains, with each domain containing between one to seven communities. This suggests the presence of dynamically or functionally related substructures within domains that exhibit slightly distinct responses to force.

**Table 2:** Dynamic community detection results on force-clamp MD datasets (*t*_1_ to *t*_4_ corresponded to the different end-to-end extension levels applied in the force-clamp simulations of *α*_5_*β*_1_: 12nm, 14nm, 16nm, 18nm; *α*_*V*_*β*_3_: 3nm, 11nm, 16nm, 18nm).

| Model | Extension Level | $\alpha_5\beta_1$ | | $\alpha_V\beta_3$ | |
| --- | --- | --- | --- | --- | --- |
|  |  | Modularity | Conductance | Modularity | Conductance |
| DEC | $t_1$ | 0.839 (0.006) | 0.131 (0.007) | 0.837 (0.003) | 0.133 (0.003) |
| DMON |  | 0.873 (0.006) | 0.086 (0.006) | 0.875 (0.002) | 0.080 (0.003) |
| DynAERNN |  | 0.189 (0.014) | 0.776 (0.015) | 0.136 (0.010) | 0.835 (0.012) |
| MFC-TopoReg |  | 0.645 (0.021) | 0.277 (0.020) | 0.628 (0.019) | 0.327 (0.018) |
| **DynMoCo** |  | **0.901 (0.002)** | **0.058 (0.005)** | **0.892 (0.002)** | **0.067 (0.002)** |
| DEC | $t_2$ | 0.842 (0.008) | 0.127 (0.008) | 0.834 (0.007) | 0.135 (0.007) |
| DMON |  | 0.869 (0.007) | 0.008 (0.090) | 0.857 (0.012) | 0.094 (0.011) |
| DynAERNN |  | 0.193 (0.013) | 0.773 (0.013) | 0.215 (0.008) | 0.756 (0.008) |
| MFC-TopoReg |  | 0.612 (0.023) | 0.309 (0.023) | 0.628 (0.017) | 0.329 (0.017) |
| **DynMoCo** |  | **0.894 (0.001)** | **0.065 (0.006)** | **0.884 (0.002)** | **0.071 (0.002)** |
| DEC | $t_3$ | 0.843 (0.007) | 0.127 (0.007) | 0.839 (0.007) | 0.130 (0.007) |
| DMON |  | 0.866 (0.004) | 0.091 (0.004) | 0.851 (0.005) | 0.099 (0.008) |
| DynAERNN |  | 0.190 (0.011) | 0.775 (0.010) | 0.199 (0.009) | 0.771 (0.008) |
| MFC-TopoReg |  | 0.632 (0.033) | 0.291 (0.032) | 0.647 (0.024) | 0.311 (0.023) |
| **DynMoCo** |  | **0.893 (0.002)** | **0.065 (0.005)** | **0.886 (0.004)** | **0.069 (0.004)** |
| DEC | $t_4$ | 0.848 (0.005) | 0.120 (0.005) | 0.841 (0.004) | 0.129 (0.004) |
| DMON |  | 0.869 (0.004) | 0.087 (0.005) | 0.851 (0.009) | 0.100 (0.009) |
| DynAERNN |  | 0.132 (0.010) | 0.833 (0.011) | 0.244 (0.013) | 0.726 (0.013) |
| MFC-TopoReg |  | 0.602 (0.019) | 0.323 (0.019) | 0.653 (0.033) | 0.303 (0.032) |
| **DynMoCo** |  | **0.894 (0.001)** | **0.065 (0.004)** | **0.884 (0.003)** | **0.072 (0.004)** |

**Table 3:** Dynamic community detection results on force-ramp SMD datasets averaged over time.

| Model | $\alpha_V\beta_3$ | | $\alpha_{IIb}\beta_3$ | |
| --- | --- | --- | --- | --- |
|  | Modularity | Conductance | Modularity | Conductance |
| DEC | 0.826 (0.001) | 0.148 (0.001) | 0.835 (0.002) | 0.137 (0.002) |
| DMON | 0.870 (0.018) | 0.091 (0.015) | 0.881 (0.002) | 0.080 (0.002) |
| DynAERNN | 0.048 (0.024) | 0.918 (0.028) | 0.076 (0.013) | 0.890 (0.013) |
| MFC-TopoReg | 0.648 (0.019) | 0.272 (0.019) | 0.612 (0.013) | 0.270 (0.013) |
| **DynMoCo** | **0.906 (0.007)** | **0.056 (0.007)** | **0.909 (0.005)** | **0.053 (0.006)** |

#### Community visualization, validation, and characterization

Sample visualization of one community overlaid on integrin is visualized in Figure S11 and for all communities in Figure 1. Communities can be visualized dynamically with the 3D molecular structure in VMD using the custom scripts provided. Community results provide a reduced representation of the high-dimensional MD simulation reduced from tens thousands of atoms into group motions of dozens of communities (Figure 2). Details of communities identified by DynMoCo are summarized in Tables S1-S4. Our post-hoc validation analysis show that the dynamic communities identified by DynMoCo are: (i) internally relatively densely connected compared to between communities (Figures 3 and S6-8), (ii) predominantly held together by covalent bonds with a significant subset of non-covalent contacts (Figure S9), and (iii) residues within communities exhibit coordinated movement (Figure S10). Analysis details are provided in the Supplemental Materials.

**Figure 1.**
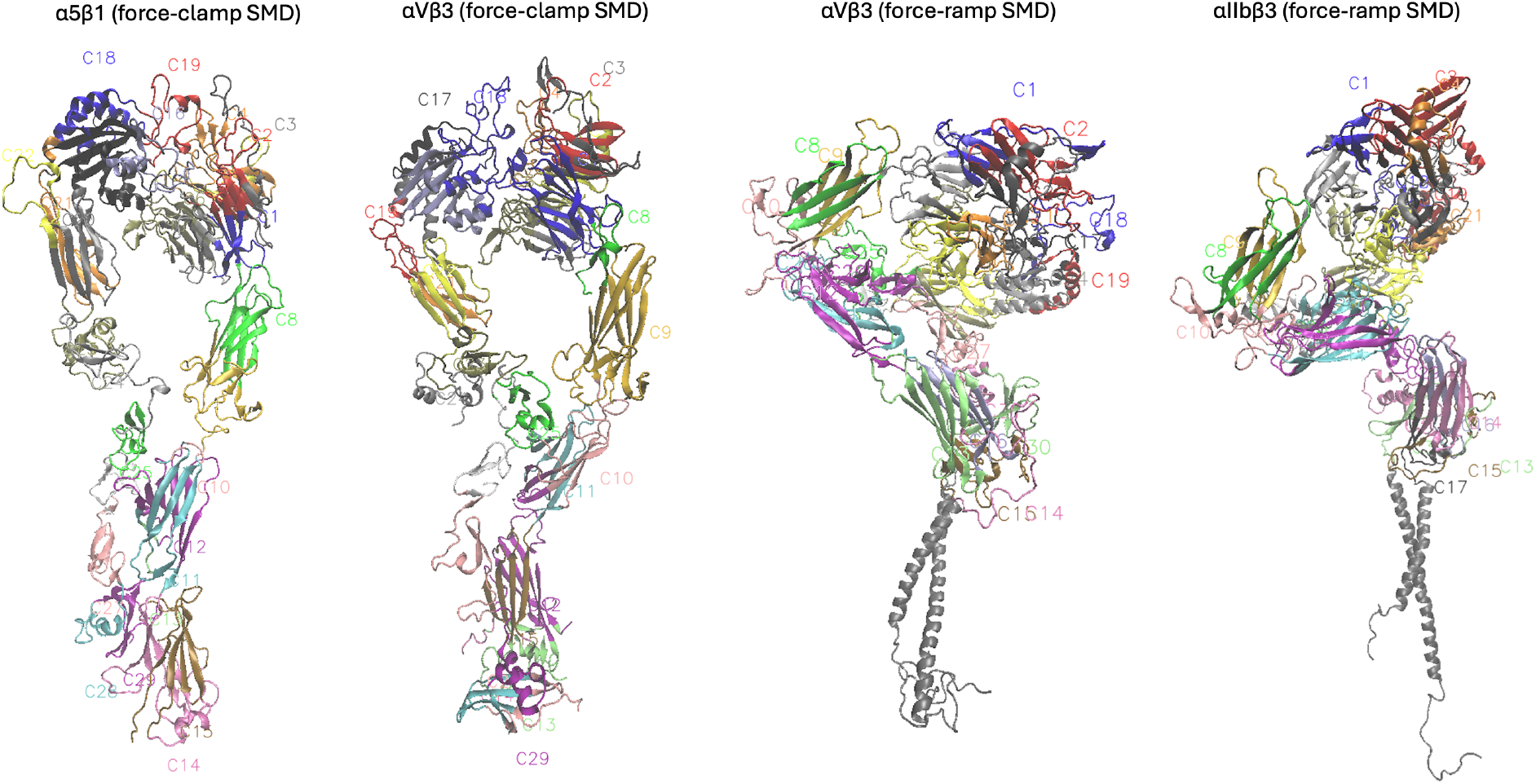
Visualization of communities overlaid on 3D molecular structure at *t*_1_. Communities are labeled as C1, C2, … 29-32 communities were identified for the three integrin systems. Communities can be visualized dynamically in VMD.

**Figure 2.**
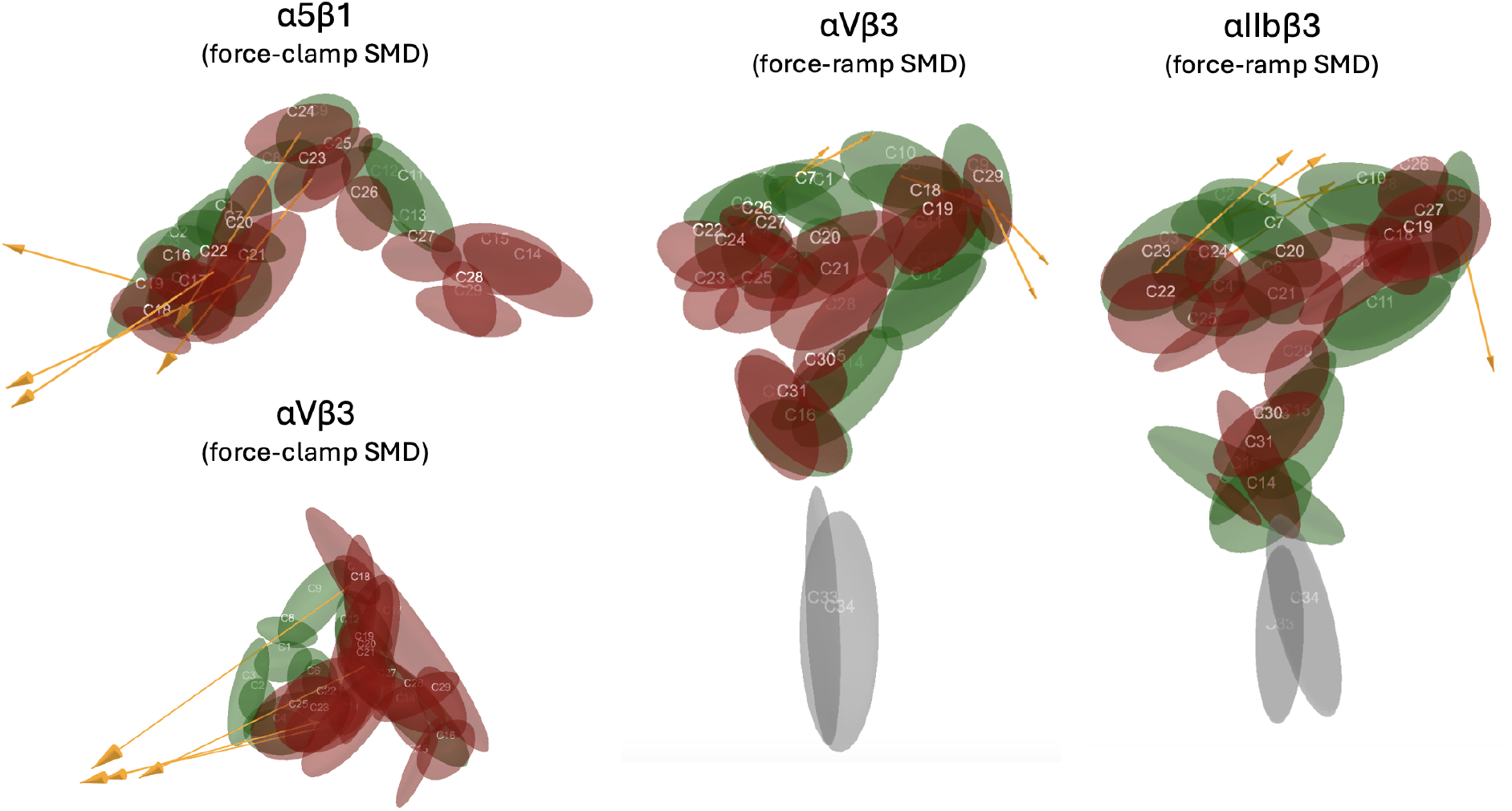
Schematics of the communities and their motion during the unbending process. The magnitude and direction of the community displacement vector from *t*_1_ to *t*_*T−*1_ is shown in yellow arrow for the top five moving communities.

**Figure 3.**
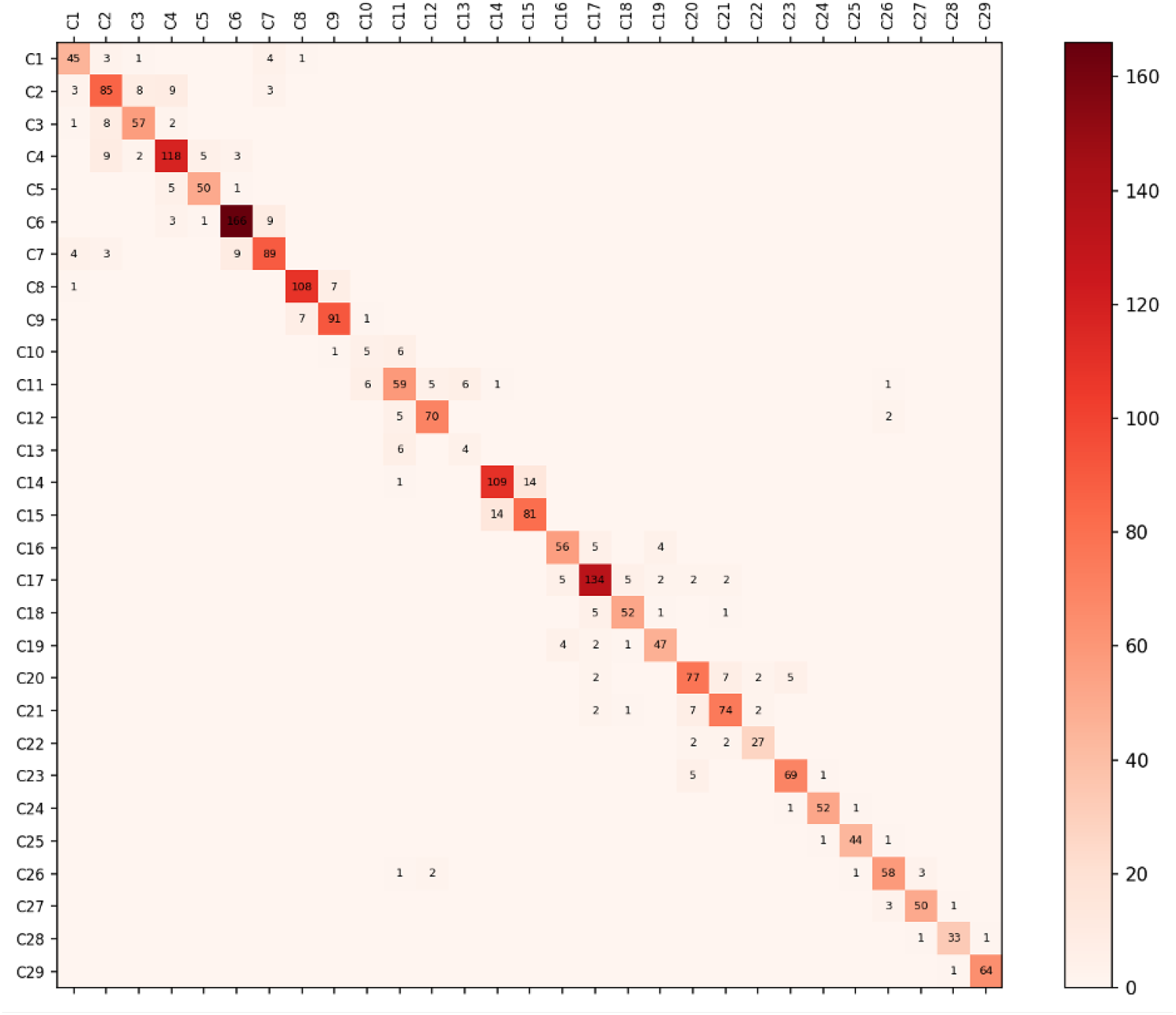
Number of covalent and non-covalent interactions within vs. across communities for *α*_5_*β*_1_ at 12nm. Communities are internally densely connected compared to connected with other communities .

#### Community structure of same integrin across different MD datasets

DynMoCo identified 29 initial communities for force-clamp *α*_*V*_*β*_3_ which are evolved into 30 communities throughout the conformational change, and 31 initial communities for force-ramp *α*_*V*_*β*_3_ which are evolved into 32 communities (Figure S2-3). While most communities identified in force-ramp and force-clamp datasets of *α*_*V*_*β*_3_ are similar, the force-ramp simulations yielded more granular communities in the thigh and *β*I domains. A major difference is that no substantial community restructuring was detected in force-ramp *α*_*V*_*β*_3_(Figure S4), whereas notable community structural changes were observed in the force-clamp simulations within the calf-1, calf-2, and *β*TD domains coupled with increases in non-covalent interactions within communities (Figure S3). The lack of community change in force-ramp simulations may stem from the force application protocol, which yielded small extension change and hence limited the observability of community evolution. Despite no community structure change was detected in force-ramp simulations, a widespread reduction in non-covalent interactions across many communities was observed, indicating relatively stable community structure coupled with local conformations internal to the communities. Both results identified communities that are mixture of residues from hybrid and *β*I domains, hybrid and PSI domains, and EGF domains, suggesting dynamic interactions of these domains in *α*_*V*_*β*_3_ during a conformational change.

#### Community structure differences across integrins

29 initial communities were identified for *α*_5_*β*_1_ at its initial state which evolved into 32 communities (Figure 4), 29 initial communities were initially identified for *α*_*V*_*β*_3_ which evolved into 30 communities (Figure S2), and 31 communities were identified for *α*_*IIb*_*β*_3_ which evolved into 32 communities (Figure S4). Overall, the community layout of the three integrins is quite alike which is expected given their structural similarity. For the two integrins with *β*_3_ chain, communities mixed with residues from hybrid and PSI domains were observed. Moreover, the community near hybrid domain showed increase in non-covalent interactions in the two integrins with *β*_3_ chain but exhibited decrease in non-covalent interactions in the integrin with *β*_1_ chain. The *α* chain was different across the three integrins and different community behaviors were observed. While no community structure change was observed in the *al pha* chain head across the three integrins, the inter-community interactions differed: *α*_5_*β*_1_’s *α* head communities showed a mixture of increase and decrease in non-covalent interactions, *α*_*IIb*_*β*_3_ exhibited a decrease in non-covalent interactions in *β*-propeller and thigh, with some of their intersection communities forming in non-covalent contacts, while *α*_*V*_*β*_3_ showed an uniform trend of either increase in non-covalent contacts in force-clamp simulations or a decreasing in non-covalent contacts in force-ramp simulations.

**Figure 4.**
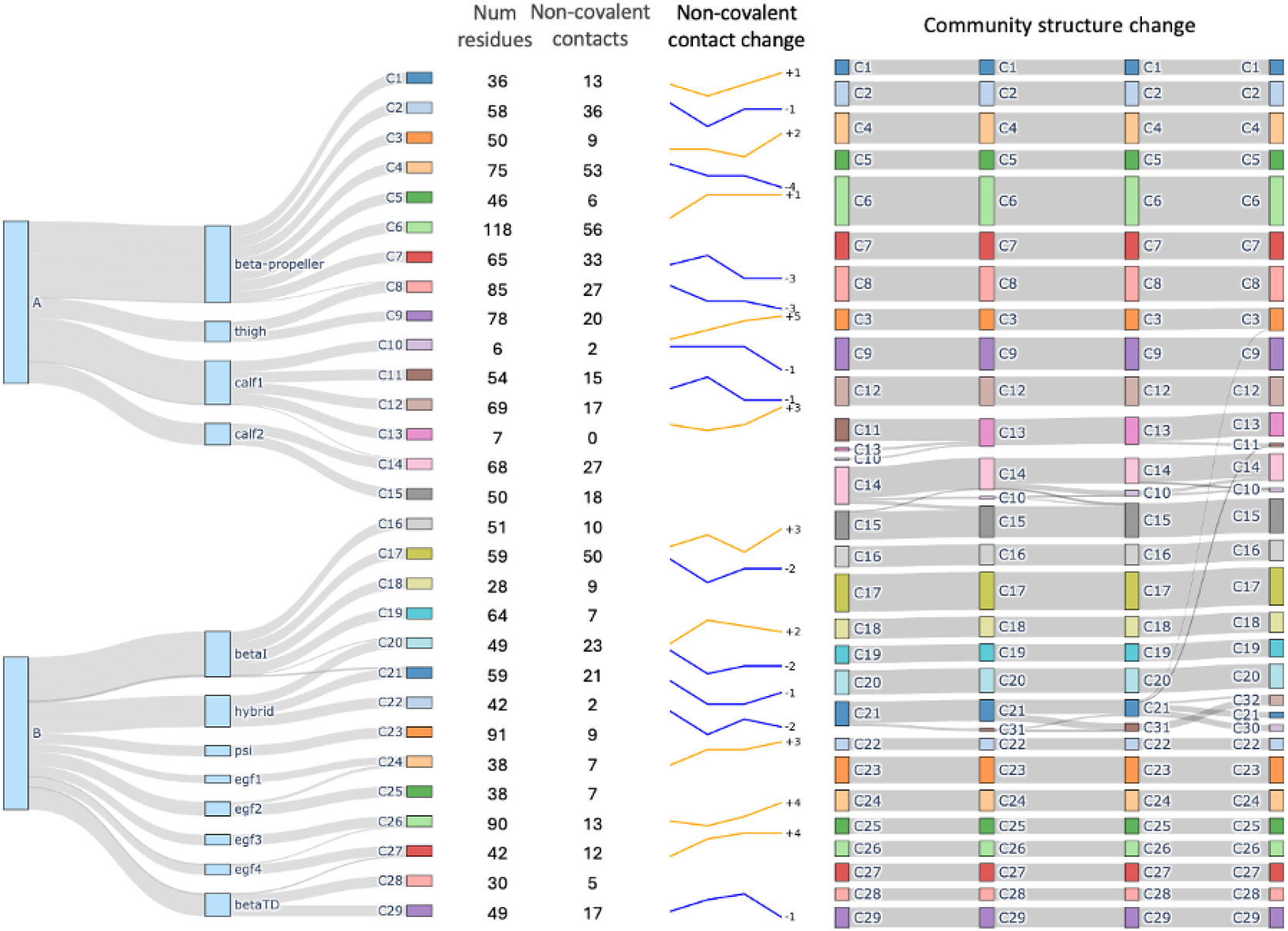
Dynamic communities detected for *α*_5_*β*_1_. Tree diagram visualizes community membership in relation to domains and chains. First column shows number of residues in each community. Second column shows number of non-covalent contacts within the community at *t*_1_. Line chart show change of non-covalent contacts over time. Last column visualize community structure change.

### Community Evolution and Conformational Dynamics

#### Community splitting and emergence in the hybrid domain for *α*_5_*β*_1_

A community (C21) identified by DynMoCo analysis of the force-clamp simulation of *α*_5_*β*_1_ drew our attention. Structurally, C21 spans the interface between the *β*I domain and the hybrid domain with member residues from both domains. Dynamically, this community displays significant temporal changes in both shape and membership during conformational changes: it began partially splitting into a new community (C31) from 12 to 14 nm, entering a split-merge state with C31 from 14 to 16 nm, and further dividing into two new communities (C30 and C32) as extension level increased to 18 nm (Figure 4). These splitting and merging events, which can be visualized in Supplemental Video S5, indicate that some residues initially showing correlated motions (i.e., belonging to C21) no longer move correlatively (i.e., changing community membership) at later times as the integrin changes its conformation. As depicted in Figure 5, which shows snapshoots of the zoomed-in movie in Supplemental Video S5(b), Thr157 (T157), a residue situated at the base of the *β*I *α*1 helix to anchor the interface between *β*I and hybrid domains, forms a network of hydrogen bonds with four other *β*I residues: Asp120 and Asp159 stay in C21 as T157 throughout the pulling simulation while Tyr121 and Met153 reside in C17 and C18. Whereas C21 contains some *β*I domain residues, many of its members reside in the hybrid domain, C17 and C18 mainly consist of residues of the hybrid domain (Figure 4). Therefore, it seems reasonable to suggest that the hydrogen network centered around T157 may provide mechanical connections between *β*I and hybrid domains. Significantly, T157 is functionally important, as the T157A mutation severely impairs fibronectin binding and prevents the exposure of activation epitopes (e.g., HUTS-4) on the hybrid domain, effectively locking the integrin in an inactive state (28). Our DynMoCo analysis suggests an explanation to this experimental finding: Eliminating these hydrogen bonds by replacing threonine with alanine at position 157 may reduce force transmission between the *β*I and hybrid domains, thereby making the hybrid domain swing-out mechanically unfavorable and preventing the integrin from changing to the open conformation necessary for effective ligand binding, as observed in the Barton et al. study (28).

**Figure 5.**
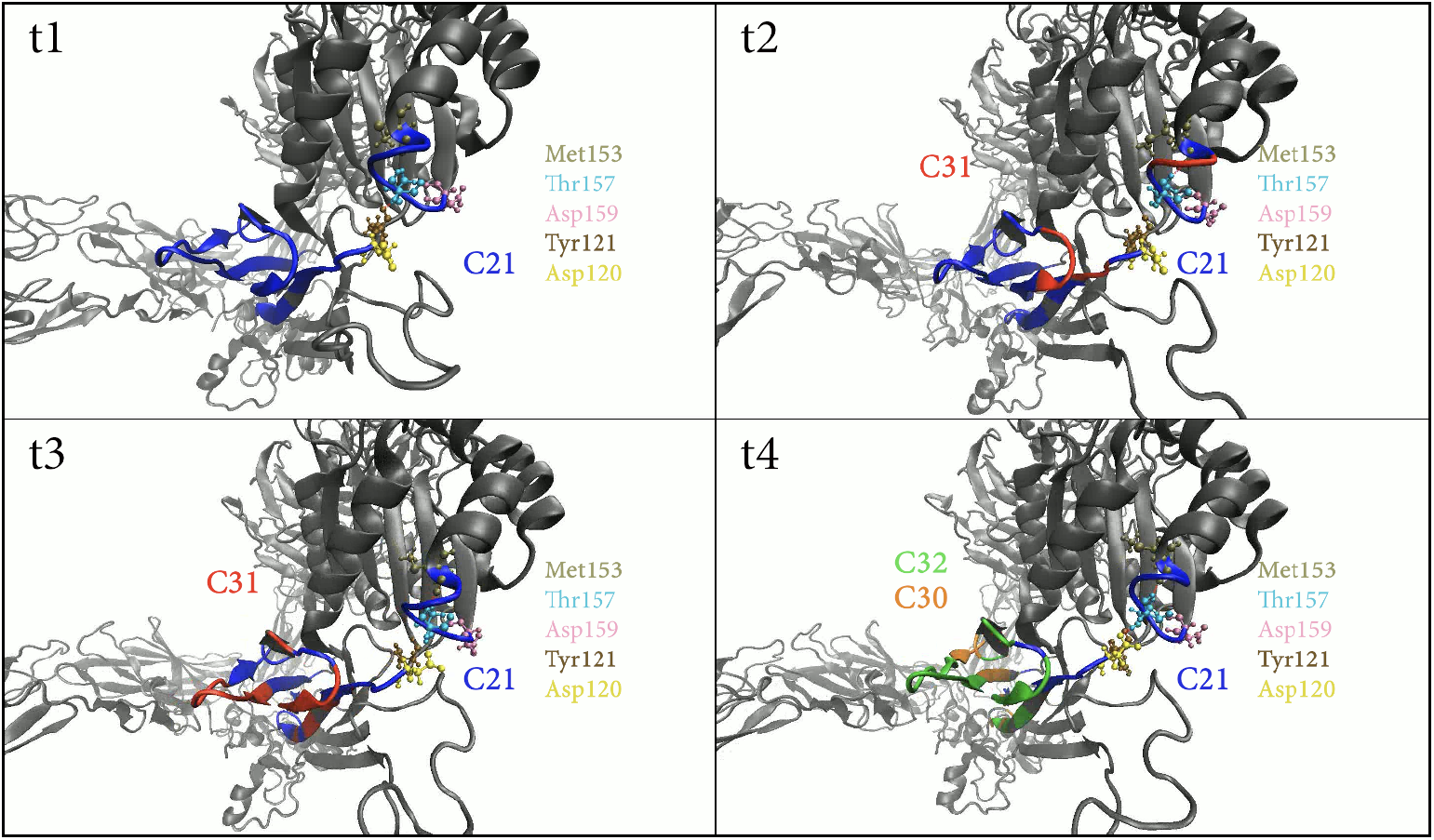
Community dynamics at the *β*_1_A–hybrid domain interface under force. Representative snapshots from the 12 nm force-clamp simulation of *α*_5_*β*_1_ integrin at frames 0, 50, 100, and 150 zoomed near the base of *α*1 helix/hybrid domain interface. The protein backbone is rendered in NewCartoon representation. The residue Thr157 is highlighted in cyan, with its primary interacting partners: Asp120 (yellow), Tyr121 (brown ochre), Met153 (tan), Asp159 (pink). Residues assigned to C21 are initially colored blue. The time-course highlights the significant dynamics of C21 as it splits into emerging communities, C30 (orange), C31 (red), and C32 (green). Visualized in VMD. See also Supplementary Video S5.

#### Community merging in the calf1-2 domain for all three integrins

In the same force-clamp simulation for *α*_5_*β*_1_, three communities (C10, C11, C13) in calf-1 domain merged into a single community during the integrin’s early extension (Figure 4). A similar phenomenon was observed in the force-clamp simulation of *α*_*V*_*β*_3_ where part of C12, C13, C14 were merged into C13 as the integrin unbent from 3nm to 11nm (Figure S2), as well as in the force-ramp simulation of *α*_*IIb*_*β*_3_, where C13, C15, C17 merged into C13 in early stages of pulling (Figure S3). These merge events were accompanied by increases in non-covalent interactions within the affected communities, suggesting tightening of local structures in the upper *α*-leg region under applied force.

#### Larger motions in communities from *β*-propeller, *β*I, hybrid, thigh domains

In the force-clamp SMD datasets, communities from the *β*-propeller, *β*I, and hybrid domains consistently exhibit the largest motions (Figure 6). The communities in the calf2 and *β*TD regions had the least motion, which is expected given its the constrained location of the pulling process during simulation. In the absence of such constraints, as in the force-ramp simulations, calf2 regions exhibited more motions (Figure 6).

**Figure 6.**
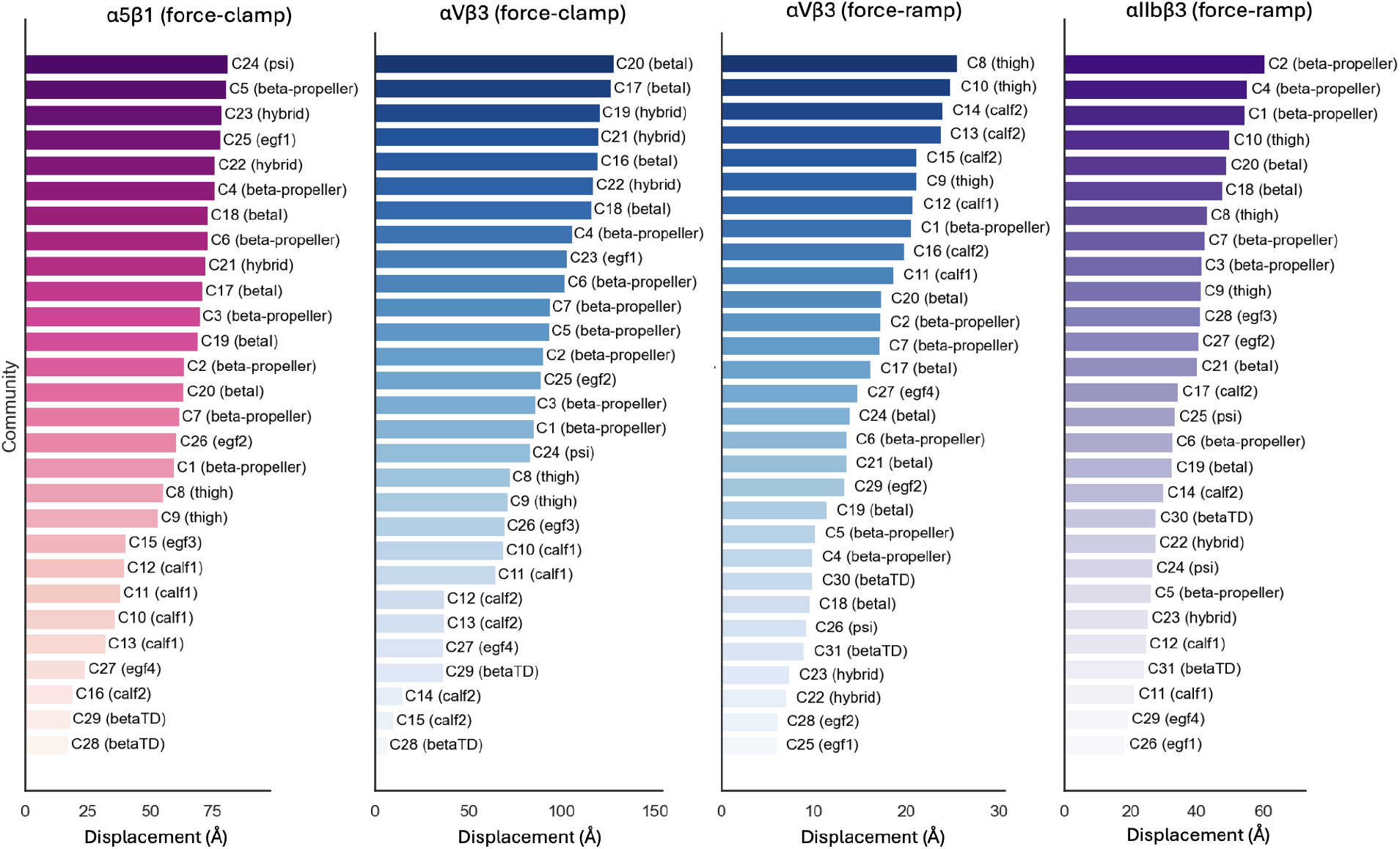
Average community displacement throughout the unbending process. Displacement is computed as the average Euclidean distance from *t*_1_ to *t*_4_ for each Ca atom and averaged across residues in each community.

A consistent trend across force-ramp SMD datasets is that communities in the *β*-propeller and thigh domains ranked among the top in displacement, the latter phenomenon was not observed in force-clamp simulations. Moreover, more rotational movement were observed in communities from the *β*TD, *β*-propeller, calf2, hybrid, and PSI regions (Figure S6), suggesting these domains undergo substantial reorientations during the force-induced transitions.

#### Greater community stability in *α*_5_*β*_1_ than *α*_*V*_*β*_3_

In force-clamp simulations, *α*_*V*_*β*_3_ underwent more pronounced and rapid structural reorganization than *α*_5_*β*_1_ (Table 4). This is reflected by its lower ARI (0.918) and Normalized Mutual Information (0.966) compared to *α*_5_*β*_1_ (ARI = 0.963, NMI = 0.979). Notably, *α*_*V*_*β*_3_ showed the greatest structural shifts early in the conformational transition (from 3nm to 11nm), indicating a quicker response to the applied force, whereas *α*5*β*1 displayed slower and evenly distributed changes throughout the trajectory. This observation aligned with previous experimental results (26) that the force-induced conformational change of *α*5*β*1 is more gradual. In force-ramp simulations, *α*_*V*_*β*_3_ and *α*_*IIb*_*β*_3_ showed on par community stability with *α*_*IIb*_*β*_3_ exhibiting slightly greater overall structural changes, particularly at the end of the extension process (Table 5).

**Table 4:** Stability of community assignments over time for *α*_5_*β*_1_ and *α*_*V*_*β*_3_ (force-clamp SMD) quantified by adjusted rand index (ARI) and normalized mutual information (NMI).

| Extension Levels | | $\alpha_5\beta_1$ | | $\alpha_V\beta_3$ | |
| --- | --- | --- | --- | --- | --- |
| Level 1 | Level 2 | ARI | NMI | ARI | NMI |
| $t_1$ | $t_2$ | 0.970 (0.008) | 0.984 (0.003) | 0.924 (0.015) | 0.971 (0.004) |
| $t_2$ | $t_3$ | 0.972 (0.008) | 0.981 (0.004) | 0.935 (0.013) | 0.974 (0.002) |
| $t_3$ | $t_4$ | 0.972 (0.009) | 0.983 (0.005) | 0.947 (0.024) | 0.976 (0.007) |
| $t_1$ | $t_4$ | 0.963 (0.015) | 0.979 (0.004) | 0.918 (0.023) | 0.966 (0.007) |

**Table 5:** Stability of community assignments over time for *α*_*V*_*β*_3_ and *αI I b β*3 (force-ramp SMD) quantified by adjusted rand index (ARI) and normalized mutual information (NMI).

| Extension Levels | | $\alpha_V\beta_3$ | | $\alpha_{IIb}\beta_3$ | |
| --- | --- | --- | --- | --- | --- |
| Level 1 | Level 2 | ARI | NMI | ARI | NMI |
| $t_1$ | $t_2$ | 0.988 (0.014) | 0.994 (0.007) | 0.985 (0.013) | 0.992 (0.006) |
| $t_2$ | $t_3$ | 0.988 (0.015) | 0.993 (0.007) | 0.991 (0.005) | 0.993 (0.003) |
| $t_3$ | $t_4$ | 0.991 (0.007) | 0.994 (0.002) | 0.987 (0.011) | 0.991 (0.006) |
| $t_1$ | $t_4$ | 0.987 (0.003) | 0.994 (0.001) | 0.973 (0.005) | 0.984 (0.003) |

### Interface Residues Relevant to Community-Community Interaction Mechanics

Beyond analyzing individual communities, it is also informative to examine *interface residues* that mediate interactions between distinct communities. We hypothesize that the nature of these interfaces influences how tightly communities are coupled in their movement under force. Specifically, a strong interface may constrain communities to move cohesively, while a loose interface may permit differential motion, acting like a flexible hinge. This analysis provides insight into how interface composition contribute to the collective dynamics of molecular substructures under force.

We define interface residues as those in meaningful contact (i.e., within 5Å) with residues belonging to a different community.

To better understand the structural role of these residues, we categorize interface residue pairs into two types: (i) Adjacent pairs: residues adjacent to one another in the protein’s sequence connected by covalent polypeptide bonds, indicating structural continuity, and (ii) Non-adjacent pairs: residues separated by at least one additional residue in the protein’s polypeptide sequence connected only via non-covalent interactions, reflecting spatial rather than sequential proximity. To quantify the mechanical coupling between communities, we calculate the average angle between displacement vectors of residues in the two communities. A large inter-community displacement angle suggests that the interface permits relative motion, indicative of a loose, hinge-like connection. Conversely, a small angle implies coordinated movement, reflecting a strong mechanical linkage between the communities.

In *α*_5_*β*_1_, the angles between adjacent communities are relatively small (mean 18.3°, sd = 9.5°), indicating tight community interfaces (Table S5). By contrast, *α*_*V*_*β*_3_ exhibits larger inter-community angles (mean 20.6°, sd = 17.7°), particularly in the calf1, calf2, *β*I, hybrid, and EGF domains, suggesting looser, hinge-like connections in these regions (Table S6). This observation is consistent with our previous finding of larger conformational motions and reduced stability in this integrin (Table 4). Similarly, in force-ramp simulations, *α*_*V*_*β*_3_ shows slightly greater inter-community angles (mean 41.1°, sd = 22.3°) (Table S7) compared with *α*_*IIb*_*β*_3_ (mean 41.1°, sd = 38.9°), with the largest angles again concentrated in the *β*I, hybrid, EGF1, and EGF2 domains (Table S8). These differences in community interfaces may shape how integrins propagate force and ultimately influence their function.

The complete set of interface analysis results are provided in Supplemental Materials. This analysis can provide some insights into how structural topology and interface composition contribute to the collective dynamics of molecular substructures under force

### Identifying the Highly Mechanically Responsive Communities and Residues

Communities in the hybrid domain exhibit high mechanical responsiveness from PageRank analysis for the three integrins, *α*_5_*β*_1_, *α*_*V*_*β*_3_ (force-clamp SMD), and *α*_*IIb*_*β*_3_ (Figure 7). Communities in *β*-propeller show high mechanical responsiveness in both force-clamp and force-ramp SMD data of *α*_*V*_*β*_3_. The most mechanically responsive residues are mainly concentrated in the knee and *β*-leg region for *α*_5_*β*_1_ and *α*_*V*_*β*_3_. Several community merging events include clusters of highly mechanically responsive residues, and a subset of these residues are interface residues (Table S13). Notably, in the force-clamp simulations of *α*_*V*_*β*_3_, multiple interface residues (VAL107, GLY405, ASP109) with high PageRank scores are observed at the merging interfaces of calf-2 communities. These results indicate that mechanically responsive residues are often enriched within merging communities and, in several cases, localized at merging interfaces. A detailed list of top residues with top PageRank scores from adjacent extension levels is summarized in Supplemental Materials.

**Figure 7.**
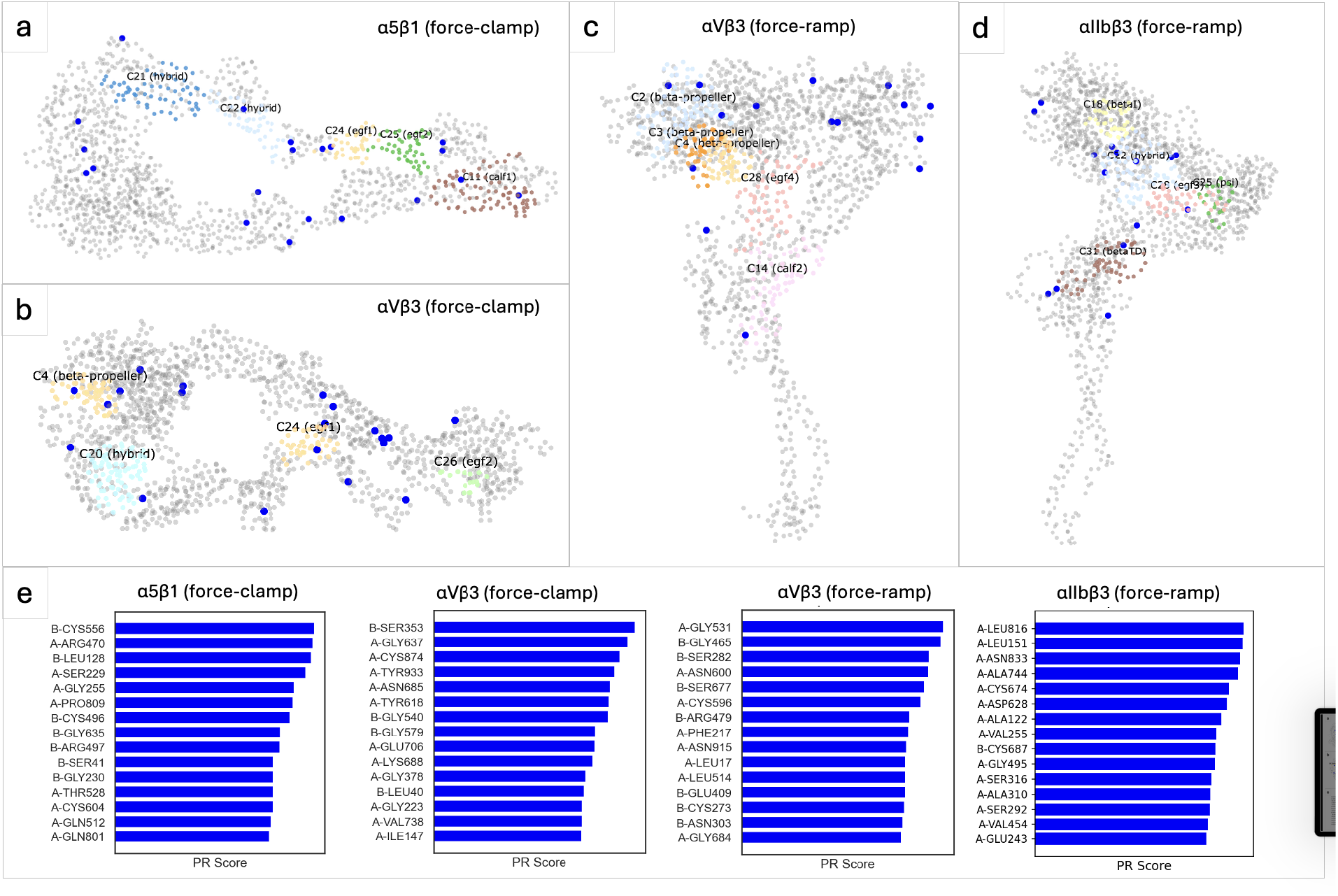
Mechanically responsive residues and communities identified by PageRank (PR). (a–d) Residue-level visualizations highlighting the top communities with the highest average PR scores (colored regions) and top residues (blue dots) identified as most mechanically responsive. (e) Bar plot showing the top 15 residues ranked by their PR scores, indicating key residues most influenced by mechanical force.

Several top mechanically responsive residues have correspondence with previous findings in the literature (Table S9-10). A-GLN189, ranked as the second highest PageRank score from 12nm to 14nm for *α*_5_*β*_1_, has been examined in a study of Rhodostomin mutants (29). A-ASP228, ranked the 7th highest PageRank score in the same analysis, was examined in a study of de novo design of *α*_5_*β*_1_-specific modulating miniprotein binders (30). Mutating the adjacent residue to A-SER156 (but not itself) was found to block the recognition of mAb 16 and perturb the high affinity binding of RGDGW-containing peptides to *α*_5_*β*_1_ (31), and this residue was found to be top mechanically responsive in the PageRank analysis of the SMD simulated transition from the 16nm to 18nm change. Another study examined mutation of B-PRO228 (32), which was identified as mechanically responsive by the analysis of the SMD simulated transtiion from 16nm to 18nm. A study on Drosophila cross spices (33) found weak associations in mutations of B-CYS374, B-GLY595, and these two residues were found to be have top PageRank scores in the analysis of the SMD simulated transitions from 3nm to 11nm as well as from 16nm to 18nm.

### Impact of *α* Chain on Community Dynamics of *β*_3_ Chain

Visual inspection of the community architectures from the force-ramp SMD simulations (Figures S3 and S4) reveals a distinct difference in the dynamic compartmentalization of the two *β*_3_ integrins. In *α*_*IIb*_*β*_3_, the dynamic community boundaries exhibit a high degree of alignment with structural domain boundaries. This indicates a modular architecture where domains move as nearly independent rigid bodies. Conversely, *α*_*V*_*β*_3_ displays significant cross-domain coupling, where single communities span multiple structural domains (e.g., C25 spans the EGF1, PSI, hybrid, and *β*-loop domains). This suggests that in *α*_*V*_*β*_3_, these distinct domains may be kinematically “locked” into larger, stiffer structural units. To translate this qualitative observation into a robust metric, we calculated the Shannon Entropy (*H*) of community membership. For a given community, *C*_*i*_, composed of residues from *m*_*i*_ domains, it is defined as

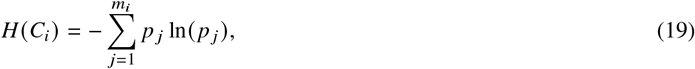

where *p* _*j*_ represents the portion of the residues in *C*_*i*_ that belong to the *j* − *th* integrin domain. This metric quantifies the diversity of structural domains contained within a single community; a value of zero indicates strict modularity (one community = one domain), while higher values indicate domain mixing.

The entropy values were calculated for *β*_3_ chain in force-ramp SMD simulations of *α*_*V*_*β*_3_ and *α*_*IIb*_*β*_3_ based on the domain composition values in Tables S5 and S7, respectively. As shown in Table S12 and Figure 8, our quantification confirms that the *β*_3_ chain in *α*_*V*_*β*_3_ exhibits >2.5x higher community entropy compared to the strictly modular profile of *α*_*IIb*_*β*_3_. This provides quantitative evidence that the mechanically-driven correlated motions of *β*_3_ chain residues are impacted by its *α*-subunit partner.

**Figure 8.**
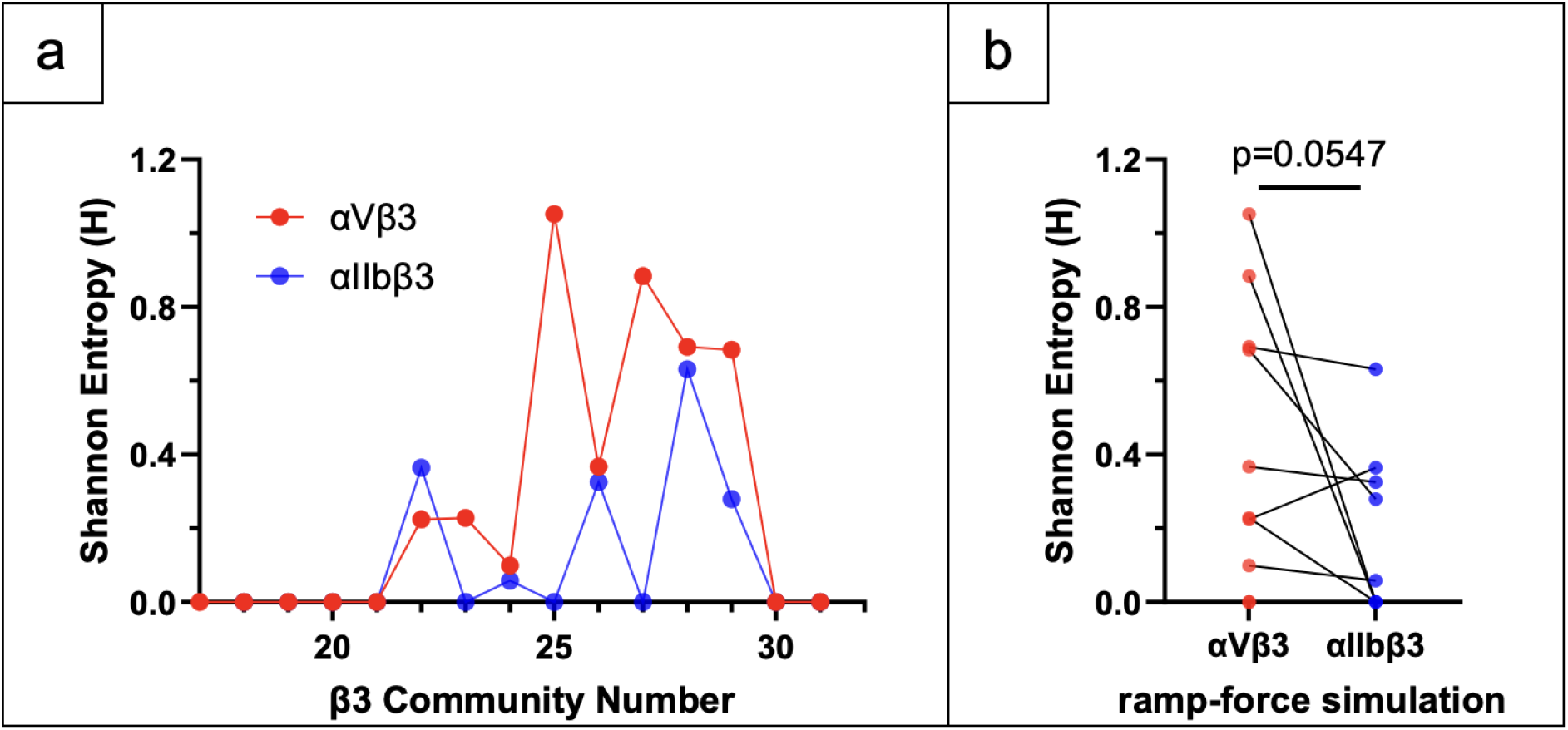
Impact of *α* chain on community membership entropy distributions of *β*_3_ chain in force-ramp SMD simulations. (a) Shannon Entropy values for each *β*_3_ community in *α*_*V*_*β*_3_ (red) and *α*_*IIb*_*β*_3_ (blue) is plotted again community number. (b) Pair-wise comparison of entropy values using the Wilcoxon matched pairs signed rank test.

These findings provide a structural mechanism for the ‘differential stiffness’ previously described in the literature (26, 27). The high entropy observed in *α*_*V*_*β*_3_ reflects a high degree of inter-domain connectivity, which likely requires higher level of motion coordination across multiple domains as opposed to rigid-body motions of individual domain relative to their hinges to achieve conformational changes. This aligns with the observation that *α*_*V*_*β*_3_ exhibits smaller variations in inter-domain distances and a ‘slower unbending’ phenotype. In contrast, the low-entropy, modular architecture of *α*_*IIb*_*β*_3_ may minimize kinematic constraints between domains to allow their relative rotation about inter-domain hinges. Since the *β*3 sequence is identical in both integrins *α*_*V*_*β*_3_ and *α*_*IIb*_*β*_3_, our results suggest that the *α*-subunit may act as an allosteric “mechanical regulator.” By altering the dynamic coupling of the *β*-subunit domains, the *α*-subunit may tailor the flexibility of the heterodimer to suit distinct physiological roles.

### Ablation Study

To evaluate the contributions of each model component of DynMoCo, we conducted an ablation study on the *α*_5_*β*_1_ dataset by systematically removing the collapse regularization, structural knowledge guidance, and temporal sparsity constraints (Table 6). The full model achieves the highest modularity (0.896) and lowest conductance (0.984), indicating strong intra-community connectivity and clear separation between communities. Removing any individual component resulted in degraded performance. In particular, eliminating the knowledge term leads to a sharp drop in modularity, underscoring the importance of domain-informed guidance for community quality. Visually examining the community structure, the full model recovers well-separated and spatially coherent communities aligned with known structural domains (Figure 9). In contrast, ablated variants without key components (e.g., knowledge guidance, collapse regularization) result in fragmented or less meaningful groupings. These results demonstrate that each component of DynMoCo contributes meaningfully to the model’s ability to detect well-structured and biologically relevant communities.

**Table 6:** Comparison of models across collapse, knowledge, sparsity, modularity, and conductance.

| Model | Collapse | Knowledge | Sparsity | Modularity | Conductance |
| --- | --- | --- | --- | --- | --- |
| Original | O | O | O | 0.896 (0.003) | 0.070 (0.003) |
| I | O | X | O | 0.694 (0.008) | 0.159 (0.010) |
| II | O | O | X | 0.837 (0.011) | 0.115 (0.011) |
| III | O | X | X | 0.690 (0.018) | 0.215 (0.014) |
| IV | X | X | X | 0.627 (0.017) | 0.148 (0.016) |

**Figure 9.**
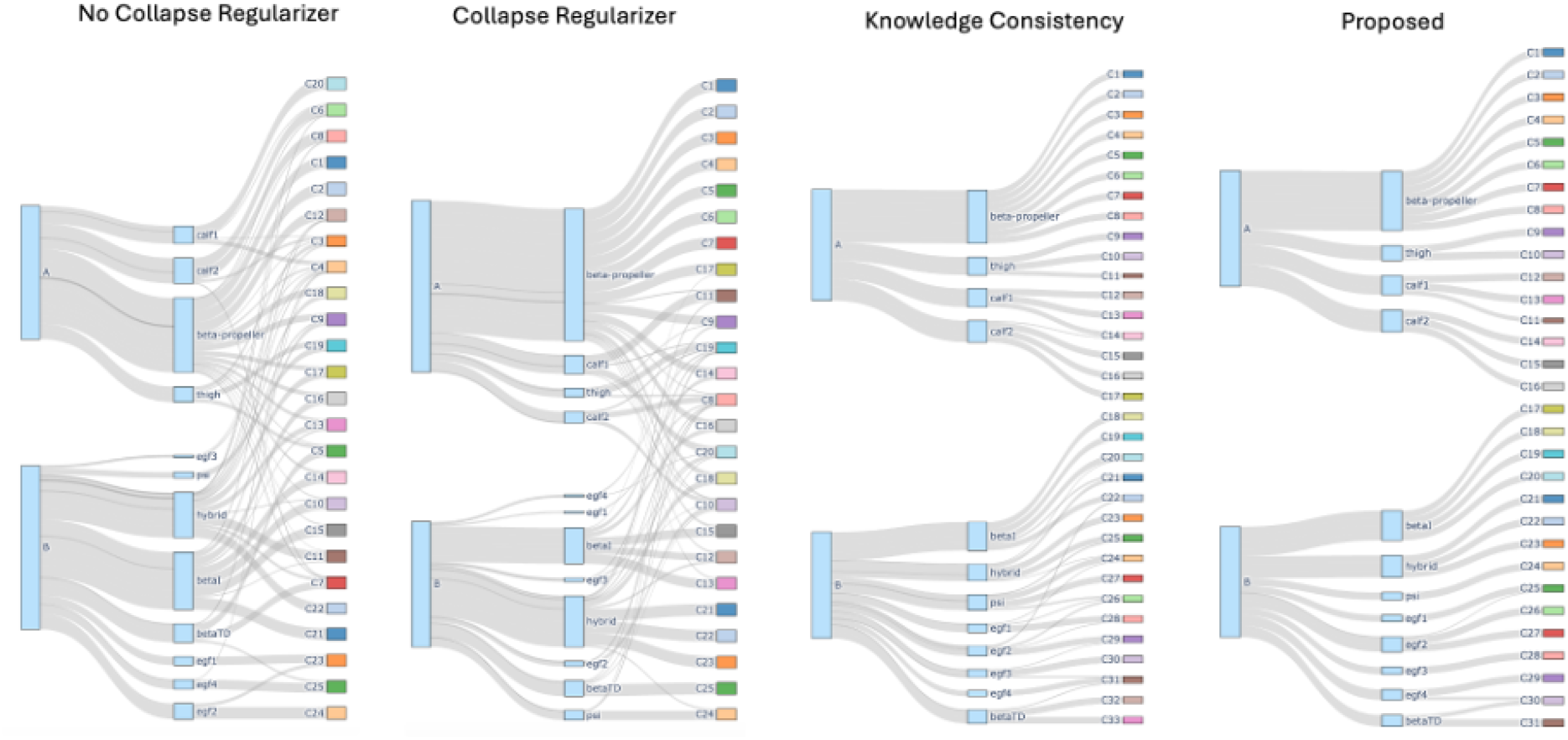
Community assignments for *α*_5_*β*_1_ using different ablated variants of the proposed DynMoCo model. The full model recovers well-separated and spatially coherent communities aligned with known structural domains. Ablated variants without key components result in fragmented groupings.

## CONCLUSION

DynMoCo developed in this work is the first end-to-end deep learning framework for dynamic community detection in molecular graphs designed for analyzing MD simulations. Methodologically, DynMoCo integrates the representational power of graph convolutional networks with the temporal modelling capability of recurrent neural networks to model dynamic graphs and identify structurally coherent and potentially functionally related communities. We applied DynMoCo to analyze force-ramp and force-clamp SMD simulations of three integrin systems simulated by two different research centers and identified around 30 communities. Characterizing the communities, analyzing community interface contacts, and complementing it with a residue-level force sensitivity analysis provided insights into the similarities and differences of the molecular dynamics of the three integrin systems. This framework allows converting high-dimensional, high-frequency 4D MD datasets into lower-dimensional, physically interpretable communities, revealing modular and local substructures within conventional domain boundaries and offering more granular insights into conformational dynamics of the molecular system. To facilitate adoption, we open-source the model code and provide a library of custom-written scripts that allow users to interactively visualize community outputs of the algorithm in VMD (19). The framework is broadly applicable across different types of SMD data, adaptable to variations in force application, restraint conditions, and molecular systems.

One limitation of the current implementation is that non-covalent interactions are modeled uniformly using a distance-based cutoff, without distinguishing their underlying physical nature (e.g., salt bridges, hydrogen bonds, or hydrophobic contacts). While this simplification enables a general and flexible framework, it may overlook interaction-specific contributions to community structure and dynamics. In addition, the method’s performance may depend on factors such as trajectory length, frame sampling frequency, temporal interval selection, and the choice of graph construction parameters (e.g., cutoff thresholds). In settings with short trajectory and few sampled frames and limited replications, deep learning methods can have overfitting risk. Future work can extend the methodology by embedding categories of non-covalent interactions as additional edge attributes or by incorporating residue-level force measurements as node attributes. Future studies will broaden the scope of applications and experimentally validate key computational findings.

## DATA AVAILABILITY

The force-clamp SMD simulation of *α*_*V*_*β*_3_ and *α*_5_*β*_1_ are provided in our Github repo. The force-ramp all-atom MD simulation of *α*_*V*_*β*_3_ and *α*_*IIb*_*β*_3_ can be downloaded from the Github repo published by the original authors in https://github.com/tamarabidone/alphaV_vs_alphaIIB.

## AUTHOR CONTRIBUTIONS

L.M., M.G., J.L., C.Z. designed the research, Z.L. provided the raw simulation data, L.M. and M.G. analyzed the data, L.M., P.C., Z.L., C.Z. intrepreted the results, A.K., S.K. developed the VMD visualization scripts, L.M. and J.P wrote the article. J.L., C.Z. supervised and administered the project. All authors reviewed the article.

## ACKNOWLEDGMENTS

This work was supported by grants from the NIH U01CA250040, U01CA280984, R01CA243486, R01CA284604 (C.Z.), and R35GM159909 (Y.C.).

